# Targeting ROCK Signaling Mitigates Influenza Virus-Induced Fibrogenesis in Human Airway Organoids

**DOI:** 10.1101/2025.04.21.649854

**Authors:** Hussin Rothan, Ahmed Mostafa, Mahmoud Bayoumi, Chengjin Ye, Ramya Barre, Anna Allué-Guardia, Aitor Nogales, Jordi B. Torrelles, Luis Martinez-Sobrido

**Author notes:** **Contact:** Hussin Rothan, Luis Martinez-Sobrido.

## Abstract

Influenza A virus (IAV) pandemics continue to pose serious global health threats, particularly to immunocompromised individuals, children, and the elderly. IAV infections trigger inflammation and tissue damage, promoting lung fibrosis. Unraveling these mechanisms is key to preventing and treating viral-induced pulmonary fibrosis and its lasting impact on respiratory health. Despite available antivirals and vaccines, there is a lack of FDA-approved therapeutics for severe or prolonged IAV pathogenesis. We modeled infection with a recombinant highly pathogenic human A/Texas/37/2024 H5N1 (rHPh-TX H5N1) strain using human airway organoids (HAO) to investigate viral replication, innate immune response, infection-induced fibrogenesis, and therapeutic approaches. The rHPh-TX H5N1 replicated efficiently, triggering a potent interferon (IFN) response and pro-inflammatory cytokine expression in HAO. Prolonged infection led to increased fibroblast-like cells surrounding infected areas, characterized by increased alpha-smooth muscle actin (α-SMA) expression and upregulation of transforming growth factor-beta (TGF-β), which caused fibroblast activation and extracellular matrix remodeling. Fibrosis-associated markers (FN, COL1A, COL3A, MMP2, MMP9) were significantly higher than in HAO infected with a pandemic recombinant A/California/04/09 H1N1 (pH1N1). Notably, Rho-associated coiled-coil-forming protein kinase (ROCK) pathway inhibition reduced fibrogenesis, with ROCK1 inhibition proving more effective than ROCK2 inhibition. These findings highlight the potential of targeting ROCK signaling to mitigate IAV-induced lung fibrosis.

## INTRODUCTION

Influenza A virus (IAV) pandemics have historically represented significant global health challenges, characterized by widespread morbidity, high mortality, and profound social disruption[1]. These pandemics arise when novel IAV strains emerge, typically through genetic reassortment between human and animal influenza viruses, causing new strain variants to evolve with limited pre-existing immunity in the human population[2–4]. Notable IAV pandemics include the 1918 “Spanish flu,” which caused an estimated 25 to 50 million deaths worldwide, the 1957 “Asian flu,” the 1968 “Hong Kong flu,” and, more recently, the 2009 “swine flu” caused by H1N1[5–7]. The emergence of highly pathogenic human and bovine Texas H5N1 (HPh/b-TX H5N1) represents a significant zoonotic threat, with its origin linked to infected dairy cows with highly pathogenic H5N1 from the clade 2.3.4.4b[8]. These events highlight the zoonotic origins of many IAV strains and underscore the importance of vigilant surveillance at the human-animal interface.

Acute IAV infection is often revealed as a self-limiting respiratory illness, with symptoms such as fever, cough, and malaise driven by the immune response to viral replication. However, severe cases may occur, particularly in individuals with compromised immunity, pre-existing health conditions, or infections involving highly pathogenic strains such as H5N1 and H7N9. Complications can arise in some cases, including viral pneumonia, acute respiratory distress syndrome (ARDS), and multi-organ failure [7,8]. Although prolonged IAV infection is rare, it is more common in immunocompromised patients, leading to persistent viral shedding, increasing vulnerability to secondary microbial infections, and impaired tissue repair. These factors contribute to long-term complications such as lung injury and fibrosis [9, 10]. Therefore, mechanistic insights into the interplay between viral replication, immune responses, and tissue repair are essential for managing severe IAV infections and mitigating their long-term consequences.

One critical complication of severe IAV infection is lung fibrosis, a debilitating and often irreversible condition characterized by excessive extracellular matrix (ECM) deposition and impaired lung function [11]. While most individuals recover from IAV infection without lasting damage, infections with highly pathogenic influenza strains or severe disease can disrupt the epithelial tissue architecture, induce chronic inflammation, and activate profibrotic pathways, such as transforming growth factor-beta (TGF-β) signaling [12, 13]. Furthermore, persistent epithelial injury, fibroblast activation, and immune cell recruitment further exacerbate fibrotic remodeling [14]. Therefore, investigating these pathological processes is crucial for identifying therapeutic targets and preventing long-term pulmonary dysfunction following IAV infections.

Human airway organoids (HAO) are being used to assess the spillover risk of highly pathogenic avian influenza (HPAI) H5N6 and H5N8 isolates [15]. This model has also emerged as a powerful system for investigating lung fibrosis mechanisms and exploring potential therapies[16]. Lung organoids possess the key features of lung architecture for studying the fibrotic changes after exposure to pro-fibrotic stimuli like TGF-β, extracellular matrix remodeling, and fibrogenic signaling pathways[17, 18]. While current limitations include incomplete immune system integration and challenges with long-term maintenance, lung organoids represent a transformative tool for understanding lung fibrosis and accelerating therapeutic discovery. Therapeutic strategies for virus-induced lung fibrosis focus on mitigating tissue remodeling and preserving lung function. Antiviral treatments may indirectly reduce fibrosis by suppressing the inflammatory response initiated by viral replication. The drug treatment programs for virus infection-related lung fibrosis are currently formulated based on the relevant guidelines for idiopathic pulmonary fibrosis (IPF), although there is no clear drug treatment program recommendation.

In this context, Rho-associated coiled-coil-forming protein kinase (consisting of two isoforms, ROCK1 and ROCK2) plays critical roles in many cellular responses to injury [19]. In this study, we utilized HAO to model infection with a highly pathogenic recombinant A/Texas/37/2024 H5N1 (rHPh-TX H5N1) strain, investigating its effects on viral replication, innate immune responses, and the development of infection-associated lung fibrosis. The virus demonstrated efficient replication within the HAO, triggering a potent interferon (IFN)-mediated antiviral response and elevated expression of pro-inflammatory cytokines. Prolonged infection with rHPh-TX H5N1 caused myofibroblast differentiation, marked by increased expression of alpha-smooth muscle actin (α-SMA). Additionally, rHPh-TXH5N1 infection was associated with the upregulation of TGF-β, which facilitated fibroblast activation and extracellular matrix remodeling. Notably, inhibition of the ROCK pathway attenuated α-SMA expression, reduced myofibroblast differentiation, and suppressed extracellular matrix protein expression, highlighting its potential as a therapeutic target to mitigate IAV-related lung injury and fibrosis. Altogether, rHPh-TXH5N1 infection induces a robust inflammatory and fibrotic response, driven by NF-κB and TGF-β signaling, which promotes airway remodeling and fibrosis. Targeting ROCK1 may offer a potential strategy to mitigate IAV-induced lung fibrosis.

## METHODS

### Biosafety

All pH1N1 or HPhTX H5N1 experiments were conducted at the Texas Biomedical Research Institute under biosafety level 2 (BSL-2) or BSL-3, respectively, and were approved by the institutional biosafety committee (IBC).

### Cell lines and compounds

Commercial human lung adenocarcinoma epithelial (A549, ATCC CCL185), and Madin-Darby canine kidney (MDCK, ATCC CCL-34) cells were maintained in Dulbecco’s modified Eagle medium (DMEM) (Invitrogen, Carlsbad, CA, USA) supplemented with 10% fetal bovine serum (FBS) and 1% PSG (penicillin, 100 U/mL; streptomycin 100 μg/mL; L-glutamine, 2 mM) at 37°C in a humidified 5% CO_2_ incubator. GSK269962A (GSK 269962), the ROCK1 selective inhibitor (CAS No: 850664-21-0), and Belumosudil (KD025; SLx-2119), the ROCK2 selective inhibitor (CAS No: 911417-87-3), were purchased from MedChemExpress (NJ, USA).

### Viruses

Recombinant pandemic A/California/04/09 H1N1 (pH1N1) and highly pathogenic human A/Texas/37/2024 H5N1 (rHPh-TX H5N1) non-structural (NS) viral segments, with the C-terminus of the non-structural 1 (NS1) protein fused either to the mCherry or Venus fluorescent proteins were used to generate pH1N1-mCherry and rHPh-TX-Venus, respectively, as previously described [20]. Briefly, NS segments were synthesized *de novo* (Bio Basic, USA) with the appropriate restriction sites for subcloning into the ambisense pDZ (pH1N1) and pHW2000 (rHPh-TX H5N1) plasmids. Modified NS segments contained the NS1 protein open reading frame (ORF) without stop codons or splice acceptor sites, followed by the porcine teschovirus-1 (PTV-1) 2A autoproteolytic cleavage site (ATNFSLLKQAGDVEENPGP), and the entire ORF of the nuclear export protein (NEP). mCherry and Venus ORFs were cloned using AgeI and NheI restriction sites, into the pDZ or pHW2000 plasmids to generate the pDZ_H1N1_NS-mCherry and pHW_H5N1_NS-Venus for virus rescues. Plasmid constructs were confirmed by complete DNA sequencing (Plasmidsaurus). Recombinant pH1N1 or HPh-TX H5N1 containing wild-type (WT) NS1 or modified NS1-mCherry or NS1-Venus, respectively, were rescued as previously described [21]. Viruses were aliquoted and stored at −80°C until use. Viral titers were calculated using the standard plaque assay in MDCK cells.

### Human airway organoid (HAO) infection

Human-derived tracheal/bronchial epithelial cells cultured at an air-liquid interface (ALI) were obtained from MatTek Life Sciences (Ashland, MA). These multilayered HAO composed of ciliated, goblet, basal, and club cells, maintained in provided medium that was changed every 3 days, were cultured into inserts in 6-well tissue plates. The fully differentiated HAO model closely mimics the epithelial architecture of the respiratory tract, exhibiting mucociliary activity and a polarized epithelium representative of the upper respiratory system. Before infection, the apical surfaces of HAO were washed twice with PBS to remove accumulated mucus. ALI cultures were infected apically with the influenza virus at a multiplicity of infection (MOI) of 0.01 to 0.001, diluted in the provided assay media. The viral inoculum was incubated on the apical surface for 2 h at 37°C, 5% CO_2_. Following this infection period, the viral inoculum was removed, and the apical surface was washed three times with PBS. Cultures were maintained at 37°C, 5% CO_2_, and viral replication was monitored over time under fluorescence microscopy (EVOS M 5000, Invitrogen). At designated post-infection times, apical washes were collected by incubating the apical surface with 200 µL of warm PBS for 30 min at 37°C, followed by collection and storage at −80°C for later viral titration. Basolateral media were also collected for cytokine and chemokine analyses.

### Virus quantification by plaque assay

MDCK cells were seeded in 6-well plates (10⁶ cells/well) and incubated overnight at 37°C in a 5% CO₂ humidified incubator. Confluent monolayers were infected with 10-fold serial viral dilutions for 1 h at 37°C. After viral absorption, cell monolayers were overlaid with agar-containing infection medium and incubated at 37°C, 5% CO₂. After 3 days, cells were fixed overnight with 10% neutral buffered formalin. To visualize viral plaques, wells were stained with 1% crystal violet for 5 min at room temperature (RT) and rinsed with tap water.

### Virus replication kinetics

The secreted mucus from HAO tissues was washed out, and the tissue was infected with rHPh-TX H5N1 in triplicate at MOI of 0.01 for 2h. After viral infection, the accumulated mucus was collected from the apical part of the insert, and the presence of infectious virus at 24-, 48-, and 72-hours post-infection (hpi) was determined by plaque assay in MDCK cells. Viral RNA was extracted from HAO tissues using Trizol and quantified by qRT-PCR. A549 cell monolayers were seeded in 6-well plates (10⁶ cells/well, triplicate) and incubated for 24 h at 37°C in a humidified 5% CO₂ incubator. Cells were then infected with the indicated viruses at an MOI of 0.001 and incubated for 1 h to allow viral adsorption. After adsorption, the virus inoculum was removed, and cells were washed three times with PBS to eliminate non-adsorbed viral particles. Fresh infection medium (3 mL) was added to each well and plates were incubated at 37°C in a humidified 5% CO₂ incubator. Cell culture supernatants (200 µL per sample) were collected at 24-, 48-, and 72-hpi and replaced with an equal volume of fresh infection medium. Viral titers were calculated using standard plaque assay in MDCK cells.

### Immunostaining

HAO tissue was washed with warm PBS and fixed in 10% formalin. Following blocking to reduce non-specific binding, tissues were incubated with a primary antibody specific to α-SMA conjugated with Alexa Fluor™ 488 overnight (Invitrogen, Cat #: 53-9760-82). Residual α-SMA antibody was washed with PBS, and the tissue was incubated for 15 min with a deep red cell mask (Invitrogen, Cat #: H32721) for cell shape visualization using fluorescence microscopy for imaging.

### RNA Extraction, cDNA Synthesis, and qPCR Analysis

Total RNA was extracted from the HAO and A549 cells using TRIzol reagent (Invitrogen, Cat #: 15596026) following the manufacturer’s instructions. Genomic DNA was removed using the DNase kit (Invitrogen, Cat. #:18068015), and RNA quantity and quality were assessed using a nanodrop. According to the manufacturer’s protocol, the cDNA was synthesized using 1 µg of total RNA using an oligo(dT) primer and reverse transcriptase kit (Invitrogen, Cat. #: 4368814). Quantitative real-time PCR (qPCR) was performed using SYBR Green Master Mix (Applied Biosystems, Cat. #: 43-091-55) on an ABI 7500 system. The thermal cycling conditions included 45 cycles of 95°C for 10 s, 60°C for 10 s, and 72°C for 10 s. Relative gene expression was analyzed using the 2⁻ΔΔCt method, with GAPDH as an internal reference gene. Primers used for amplification of genes are shown in **Table 1**.

**Table 1:**
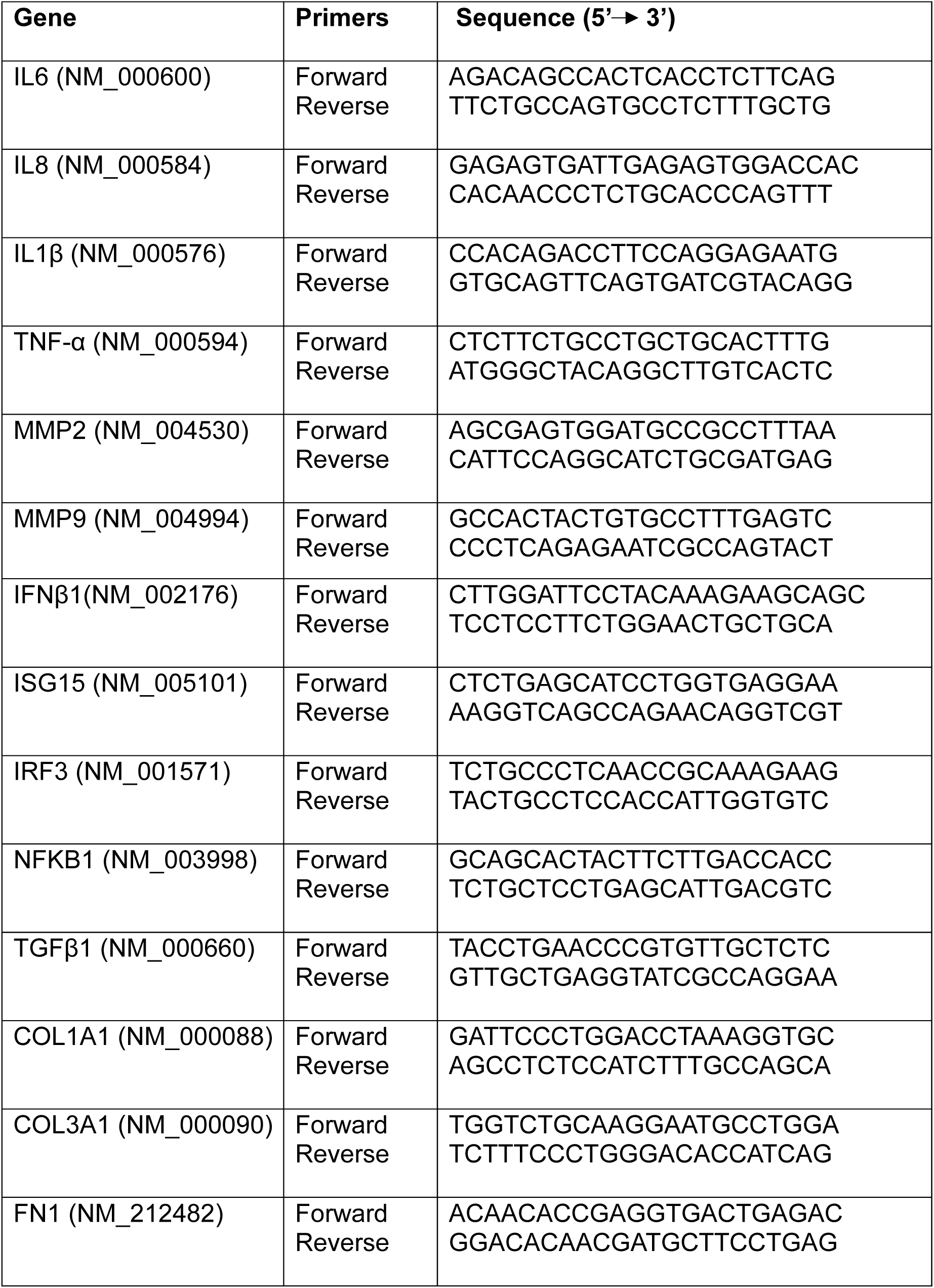
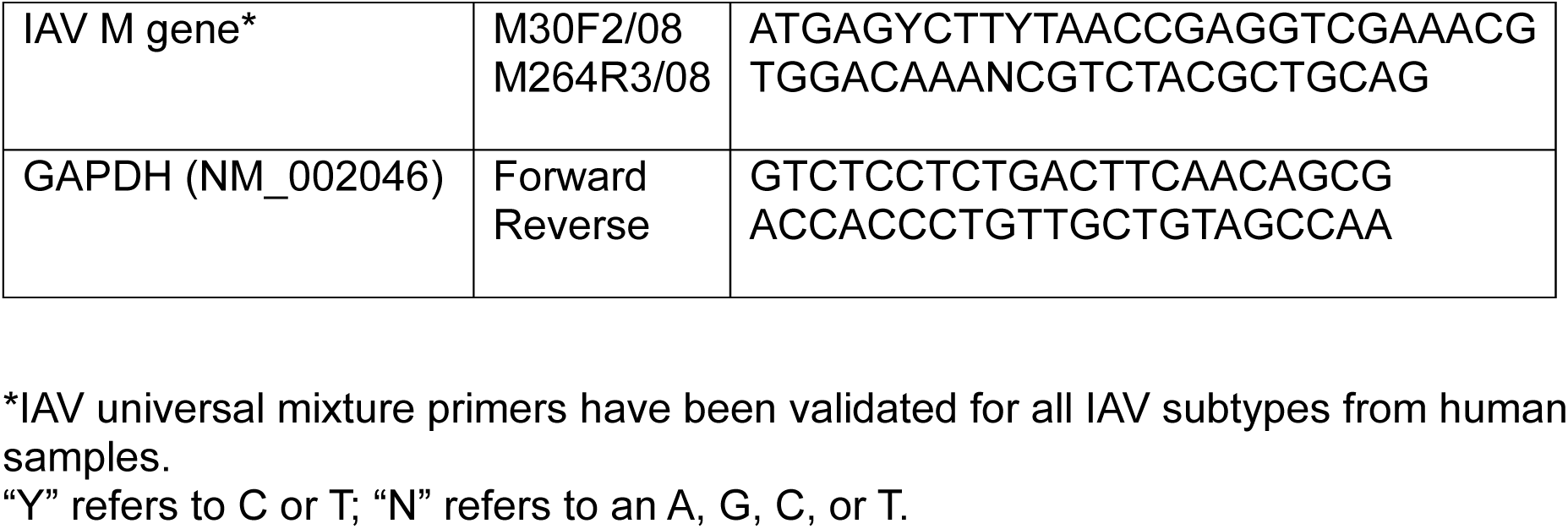
Primers used in this study.

### Multiplex Immunoassay

Levels of cytokines (IL-6, IL-1β, TNF-α) and chemokines (IP-10/CXCL-10, RANTES/CCL5) were measured using a custom human ProcartaPlex panel (Thermo Fisher Scientific, Cat. #: PPX-12-D5619750), following the manufacturer’s protocol. Mucus samples and basal media of HAO were collected at 48 hpi with rHPh-TX H5N1. Samples were centrifuged at 10,000 × *g* for 5 min to remove debris and diluted 1:2 in Universal Assay Buffer before analysis. The assay was conducted under BSL-3 conditions, and samples were decontaminated by overnight incubation in 1% formaldehyde before readout on a Luminex 100/200 System (xPONENT v4.3.309.1). Instrument settings included a gate range of 7,500–25,000, a 50 µL sample volume, 50 events per bead, a 60 s sample timeout, and standard PMT settings. Data were analyzed using xPONENT v4.3.309.1.

### Statistical analysis

Data were analyzed using GraphPad Prism 9.5.1 software. Viral titers, cytokines, chemokines, and fibrosis markers levels were compared using *t*-test, one-way, or two-way ANOVA with post-hoc corrections as appropriate. *P*-values <0.05 were considered statistically significant.

## RESULTS

### rHPh-TX H5N1 replicated efficiency in HAO

Human airway epithelial tissue is the primary site of IAV infection and replication. The air-liquid interface model of HAO (**Fig.1A**) has been considered a platform to assess viral tropism and infectivity in humans for pandemic risk assessment of zoonotic IAVs [22]. We infected HAO with rHPh-TX H5N1 at MOI 0.01 to examine virus replication. Virus shedding that occurred apically in the secreted mucus was measured via titration in MDCK cells by plaque assay, while viral RNA in the epithelial tissue was quantified using qRT-PCR. The rHPh-TX H5N1 exhibited higher replication efficiency in HAO at 48 hpi measured by the levels of infectious particles in the secreted mucus (**Fig. 1B**) and viral RNA levels in infected tissues (**Fig. 1C**). For comparison, we infected A549 cells with rHPh-TX H5N1 at MOI 0.01. Similarly, the infectious virus titers in the culture supernatants and viral RNA levels in cell lysates from infected A549 cells were higher at 48 hpi (**Figs. 1D and 1 E, respectively**).

**Figure 1:**
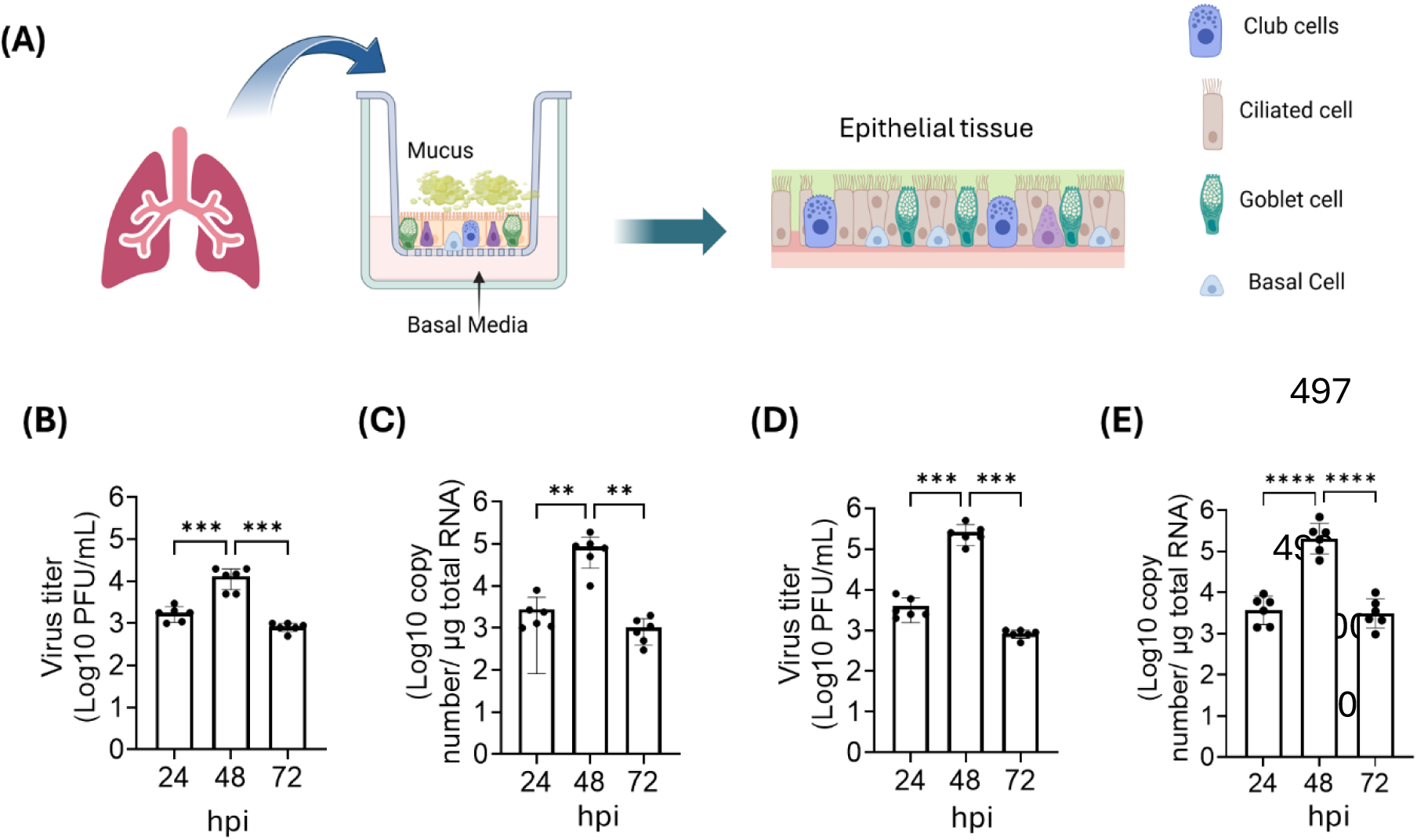
Replication kinetics of rHPh-TX H5N1 in HAO. **(A)** A diagram shows human-derived tracheal/bronchial epithelial cells cultured at an ALI. The HAO, composed of ciliated, goblet, basal, and club cells, was grown on inserts in 6-well plates. **(B)** HAO were infected in triplicate at MOI 0.01 with rHPh-TX H5N1 for 2h after washing out the secreted mucus. After infection, the accumulated mucus was collected from the apical part of the insert, and the presence of infectious virus at the indicated hours post-infection (hpi) was determined by plaque assay in MDCK cells. **(C)** HAO-infected tissues were collected for viral RNA extraction and quantified by qRT-PCR. **(D)** A549 cells were grown in 6-well plates (10⁶ cells/well, triplicate) and infected (MOI 0.01) with rHPh-TX H5N1 for 1h. Virus titers in cell culture supernatants were measured by plaque assay in MDCK cells. **(E)** qRT-PCR was used to measure viral RNA in cell lysates. Data were analyzed using one-way ANOVA followed by a post-test comparison. The significant differences are indicated (** = *p* < 0.01, *** = *p* < 0.001; **** = *p* < 0.0001; non-significant = ns).

### rHPh-TX H5N1 induces high IFN and pro-inflammatory responses in HAO

We next examined the expression levels of key players of the IFN pathway following rHPh-TX H5N1 infection of HAO. Control and infected tissue samples of HAO and A549 cell lysates were collected for RNA extraction and qRT-PCR. Expression levels of key factors of the antiviral signaling pathways were significantly upregulated upon infection with rHPh-TX H5N1 (**Fig. 2A**). For comparison, we assessed expression levels of the same factors in A549 cells after infection with rHPh-TX H5N1 (**Fig. 2B**). Generally, the IFN response of HAO to rHPh-TX H5N1 infection was lower than in A549 cells, as the HAO cell types might have varied responses representing more accurately the actual response in human airway tissues compared to the synchronized response observed in A549 cells. Indeed, the highest IFN beta (IFN-β) levels in rHPh-TX H5N1-infected HAO was observed at 48 hpi, ∼20-fold higher than mock-infected HAO (**Fig. 2A**). We observed ∼75-fold induction in infected A549 compared to mock-infected cells (**Fig. 2B**). IFN regulatory factor 3 (IRF3) and nuclear factor-kappa B (NF-κB) showed ∼2-3-fold higher expression levels in rHPh-TX H5N1-infected HAO *vs.* mock infected when compared to ∼10-15-fold observed in A549 cells (**Figs. 2A and 2B**). IRF3 and NF-κB are critical transcription factors for the induction of type I IFN and pro-inflammatory cytokines. Interestingly, NF-κB significantly increased at 72 hpi in infected HAO; however, it dramatically decreased in A549 cells (**Figs. 2A and 2B**). The upregulation of IRF3 and NF-κB led to higher IFN-stimulated gene 15 (ISG15) expression at 24 hpi in infected HAO (∼6-fold), while in A549 cells it was ∼50-fold (**Figs. 2A and 2B**). Altogether, these results demonstrate that rHPh-TX H5N1 induces a robust IFN response in HAO and A549 cells, indicating a strong activation of the host’s innate immune system.

**Figure 2:**
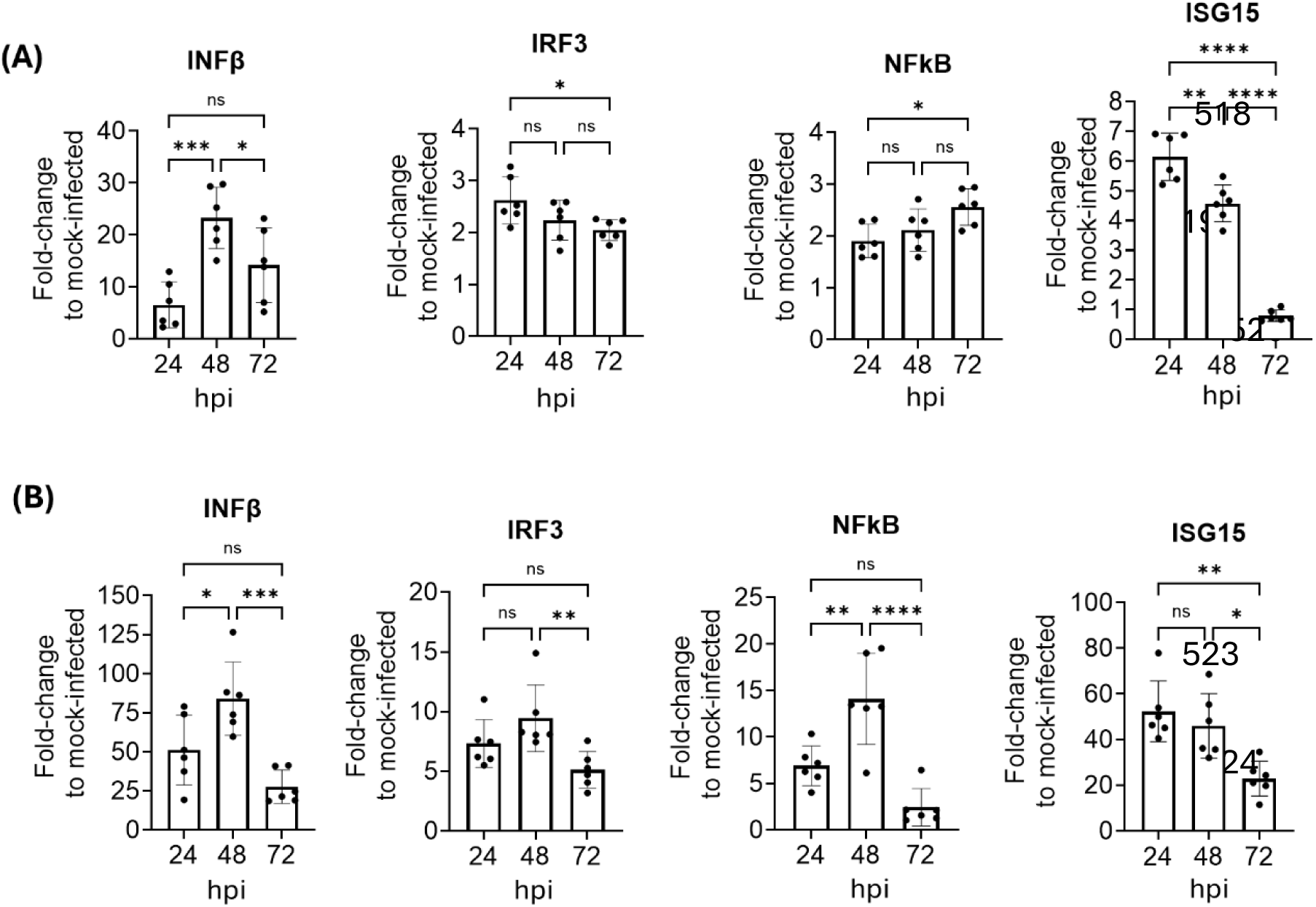
IFN responses to rHPh-TX H5N1 infection in HAO and A549 cells. **(A)** HAO were mock-infected or infected with rHPh-TX H5N1 (MOI 0.01) for 2 h, and the tissues were collected at 24, 48, and 72 hpi for RNA extraction by Trizol. Relative gene expression to mock-infected cells after normalization with the endogenous gene (GAPDH) was determined by qRT-PCR. **(B)** A549 cells were infected with rHPh-TX H5N1 (MOI 0.01) for 1 h. Mock and virus-infected A549 cells were collected at 24, 48, and 72 hpi for RNA extraction by Trizol, and gene expression levels were determined by qRT-PCR. Data were analyzed using One-way ANOVA followed by a post-test comparison. The significant differences are indicated (* = *p* < 0.05, ** = *p* < 0.01, *** = *p* < 0.001; **** = *p* < 0.0001; non-significant = ns).

To examine the inflammatory status of infected HOA, we measured pro-inflammatory markers in secreted mucus and the basal media using a multiplex cytokine assay (**Fig. 3A**). Results showed elevated levels of pro-inflammatory cytokines such as TNF-α, IL-6, IL-1β, and chemokines CCL5 (RANTES), and IP-10 (CXCL10) after infection with rHPh-TX H5N1. Higher levels of these markers were found in the HAO mucus compared to the basal media, which correlated with the viral titers observed (**Figs. 3B and 3C**). The excessive production of these mediators suggests that rHPh-TX H5N1 may trigger potent antiviral signaling pathways; however, it may also contribute to immunopathology.

**Figure 3:**
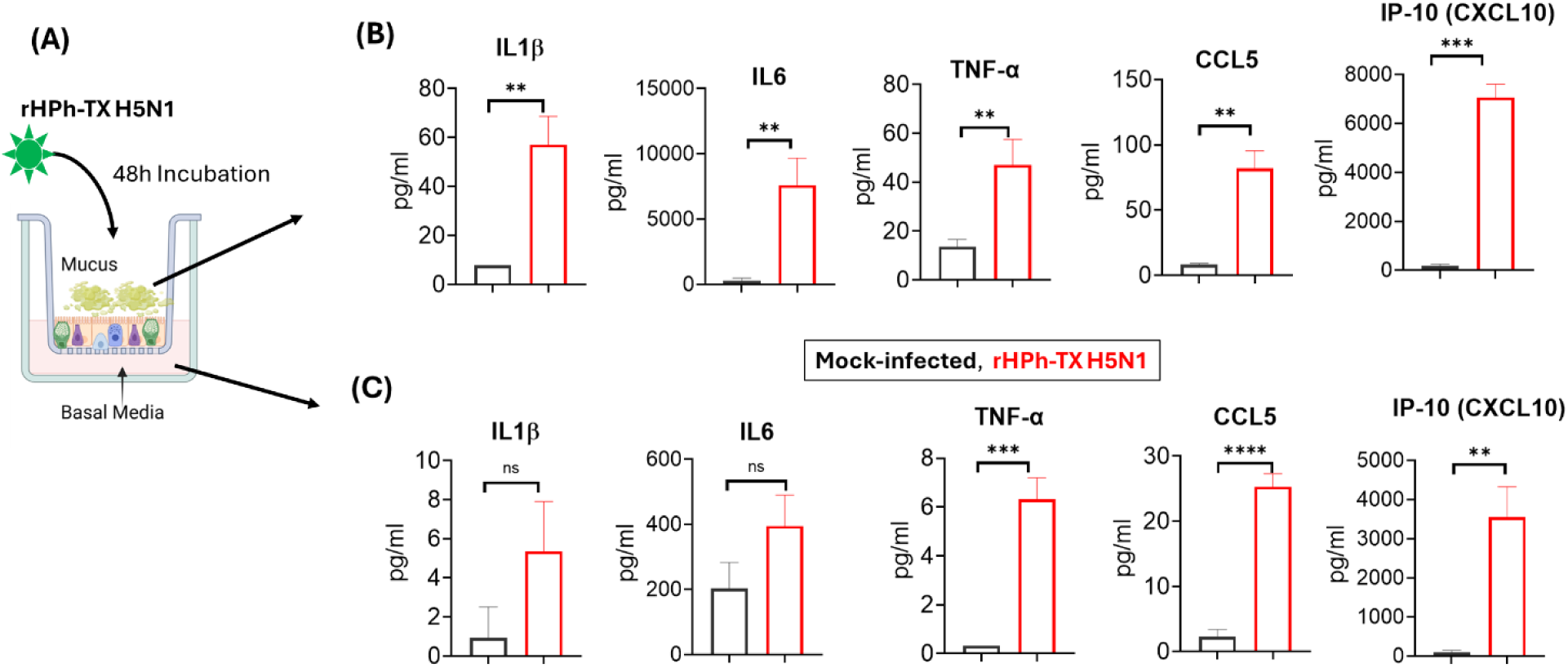
Pro-inflammatory cytokine and chemokine levels in HAO infected with rHPh-TX H5N1. **(A)** HAO inserts were infected with rHPh-TX H5N1 (MOI 0.01) for 48 h. **(B)** Secreted mucus in the apical part of the insert was collected at 48 hpi by adding 200 µL warm PBS and incubating for 15 min at 37°C. Mucus samples and **(C)** basal medium were collected at 48 hpi, spun down, and used for the multiple cytokine assay. Data were analyzed using the *t*-test comparison. The significant differences are indicated (** = *p* < 0.01, *** = *p* < 0.001; **** = *p* < 0.0001; non-significant = ns).

### rHPh-TX H5N1 and pH1N1 infections induce fibrogenesis in HAO

HAO derived from bronchial progenitor cells and differentiated to pseudostratified tissue closely mimics the morphology and function of human airways under near-physiological conditions [23]. Using this model, we next investigated the effects of rHPh-TX H5N1 infection on fibrogenesis activation compared to pH1N1 infection. Pulmonary fibrosis in pH1N1-infected patients is related to the increased activity of fibroblasts in the post-inflammatory repair pathways [24]. To monitor prolonged virus infection in HAO, we generated recombinant pandemic A/California/04/09 H1N1 expressing mCherry (pH1N1-mCherry) and recombinant A/Texas/37/2024 H5N1 expressing Venus (rHPh-TX H5N1-Venus) fused to the C-terminus of NS1 from their modified NS segments. These reporter viruses replicated efficiently in MDCK cells, achieving viral titers of ∼10^7^ PFU/ml (HPh-TX H5N1-Venus) and ∼10^6^ PFU/ml for pH1N1-mCherry (data not shown). The reporter rHPh-TX H5N1-Venus and pH1N1-mCherry were used to monitor infection dynamics in HAO over 10 days (**Figs. 4A and 4B**). Both recombinant viruses express different fluorescent proteins to eliminate any possible interaction of the reporter genes with the formation of fibroblast-like cell foci. We infected HAO with rHPh-TX H5N1-Venus and pH1N1-mCherry at an MOI of 0.001. Peak viral replication of these viruses was observed on day 2 post-infection (2-DPI), as determined by fluorescent expression, followed by a gradual decline over the subsequent days (**Figs. 4A and 4B**). By 7-DPI, consolidation areas enriched with elongated fibroblast-like cells were evident, surrounding the infected regions. These fibroblast-like cells progressively accumulated, isolating infected areas from the surrounding tissue, potentially reflecting localized host responses to infection. These structures were prominent at the latest times post-infection (e.g. 7-DPI to 10-DPI) (**Figs. 4A and 4B**). Viral titers in secreted mucus were measured by plaque assay using MDCK cells (**Fig. 4C**). The highest viral titer for both viruses was at 2-DPI, declining to undetectable levels at 7- to 10-DPI. Interestingly, viral RNA was detected at 10-DPI despite the absence of infectious virus in mucus (**Fig. 4D**). Consistently, we observed increased expression of pro-inflammatory cytokine responses in the HAO tissue harvested on 10-DPI, reflecting inflammatory status (**Figs. 4E-H**). TNF-α, IL-6, IL-8, and IL-1β showed higher levels in prolonged infection of HAO with rHPh-TX H5N1-Venus *vs.* pH1N1-mCherry (**Figs. 4E-H**). These data indicate that after 10-DPI, viral RNA can still be detected in HAO despite the lack of infectious virus in secreted mucus, with consistent increased expression of pro-inflammatory cytokines.

**Figure 4:**
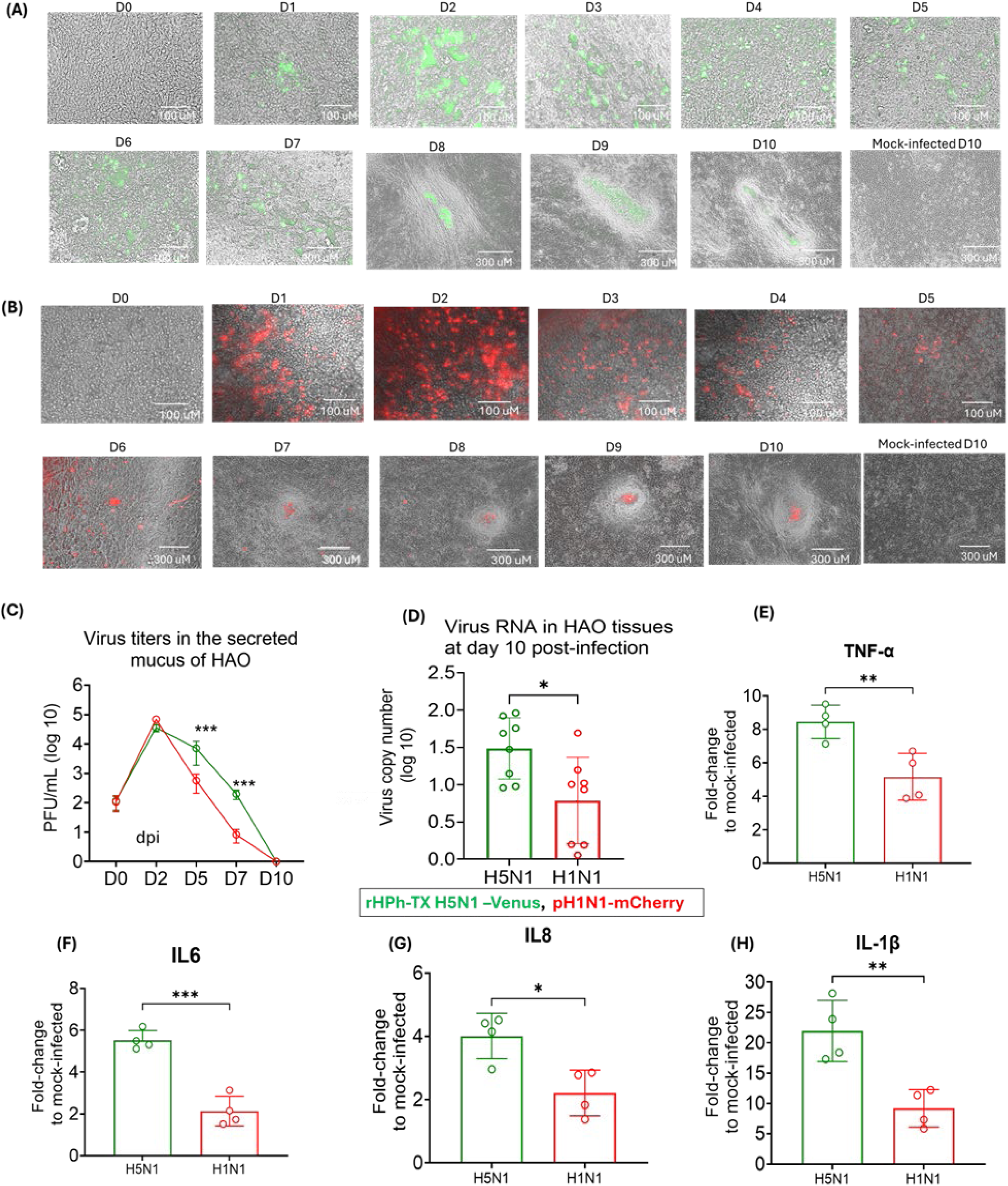
Prolonged infection of HAO with rHPh-TX H5N1-Venus and pH1N1-mCherry induces fibroblast-like cell foci surrounding infected areas and inflammation. **(A, B)** HAO inserts were infected (MOI of 0.001) with (A) rHPh-TX H5N1-Venus or (B) pH1N1-mCherry for 2 h and monitored daily under fluorescence microscopy for 10 days. **(C)** Virus titers in HAO-secreted mucus were determined by plaque assay. **(D)** On 10-DPI, virus RNA levels in HAO-infected tissues were determined by qRT-PCR. **(E-H)** Gene expression levels of pro-inflammatory cytokines in the HAO tissues at 10-DPI. Data were analyzed using two-way ANOVA and the unpaired *t*-test. The significant differences are indicated (* = *p* < 0.05, ** = *p* < 0.01, *** = *p* < 0.001; **** = *p* < 0.0001; non-significant = ns).

### rHPh-TX H5N1 and pH1N1 infections upregulated fibrogenesis markers in HAO

Our data showed alteration in the HAO epithelial tissue’s structure by forming fibroblast-like cells surrounding the virus-infected area during rHPh-TX H5N1 and pH1N1 infection. To investigate fibrogenesis activation, we next evaluate whether rHPh-TX H5N1 and pH1N1 infections induce fibroblast differentiation into myofibroblasts by α-SMA. Fluorescent staining revealed extensive FITC signal in fibroblast-like cell foci, with α-SMA filaments localized with spindle-like cell shapes, confirming the myofibroblast phenotype surrounding virus-infected cells (**Figs. 5A and 5B**). These activated fibroblasts are characterized by spindle morphology with intracytoplasmic stress fibres, a contractile phenotype, expression of various mesenchymal immunocytochemical markers like α-SMA, and collagen production [25]. The spindle morphology of fibroblasts and the high level of α-SMA expression suggest fibroblast-to-myofibroblast differentiation as part of tissue remodeling.

**Figure 5:**
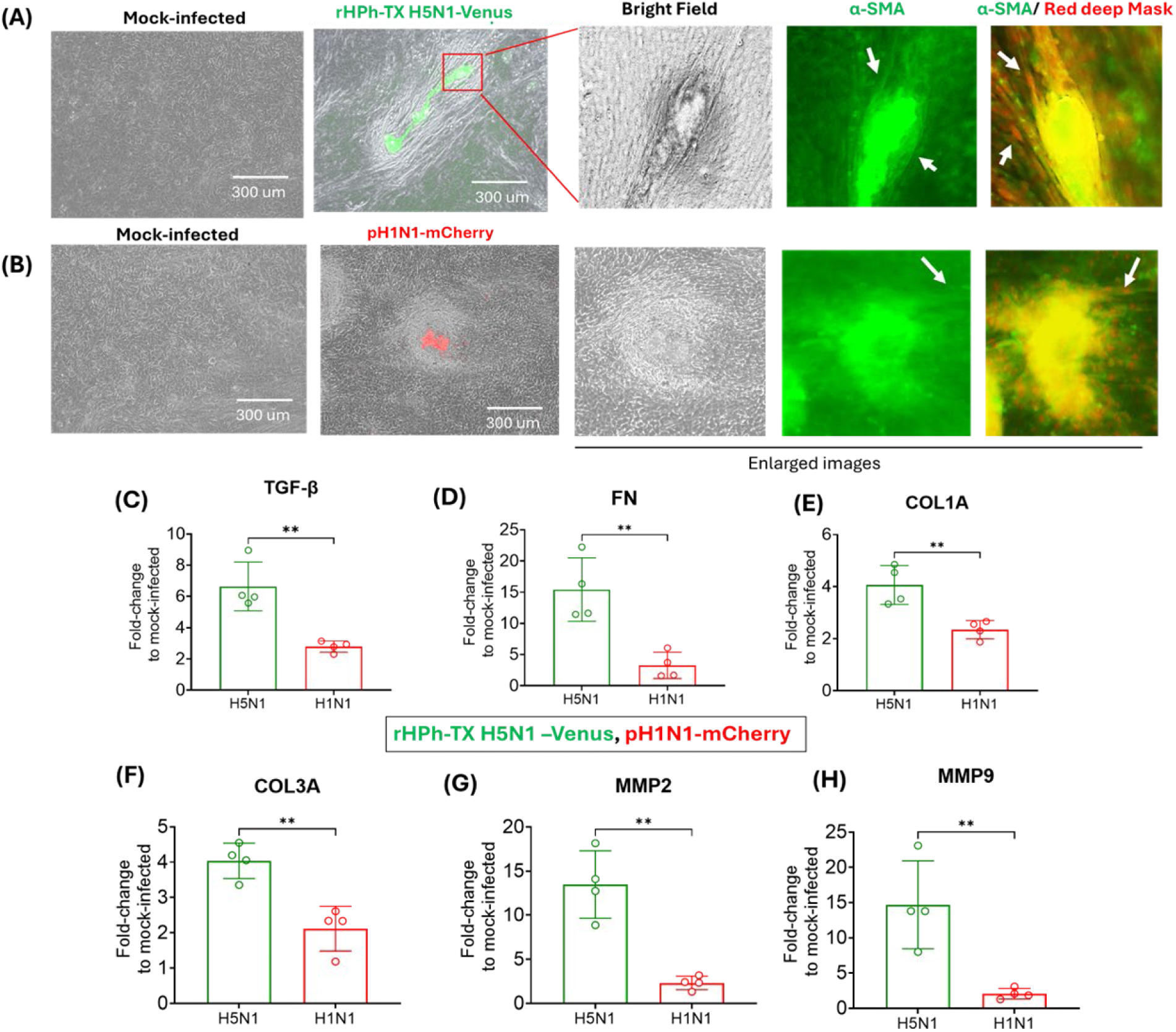
High expression of α-SMA after prolonged infection of HAO with HPh-TX H5N1-Venus and pH1N1-mCherry. **(A, B)** HAO were separately infected (MOI of 0.001) with (**A**) HPh-TX H5N1-Venus and (**B**) pH1N1-mCherry for 2 h. At 10-DPI, HAO tissue was washed 2X with PBS, fixed with 10% formaldehyde, permeabilized with 0.5% Triton-100, and incubated with Alexa Fluor 488-labelled α-SMA antibody overnight. The tissue was incubated with a red deep cell mask for 15 min to vascularize the cell shape. **(C-E)** Gene expression of TGF-β and extracellular matrix proteins fibronectin (FN), collagen 1A (COL1A), and collagen 3A (COL3A). (**F**,**G**) Gene expression tissue remodeling MMP2 and MMP9. Data were analyzed using an unpaired *t*-test, ** = *p*<0.01.

The epithelial-mesenchymal transition (EMT) has recently received consideration as a process by which epithelial cells lose cell-cell attachment, polarity, and epithelial-specific markers, undergo cytoskeletal remodeling, and gain a mesenchymal phenotype [26, 27]. TGF-β induces epithelial-mesenchymal alterations and myofibroblast transformation that contribute to lung fibrosis [19]. rHPh-TX H5N1-Venus or pH1N1-mCherry caused upregulation of TGF-β gene expression at 10-DPI, compared to mock-infected HAO (**Fig. 5C**). However, rHPh-TX H5N1-Venus induced ∼7-fold higher expression of TGF-β, while pH1N1-mCherry only induced ∼2.5-fold higher expression of TGF-β (**Fig. 5C**). Besides the role of TGF-β as a key profibrotic cytokine-promoting fibroblast proliferation, TGF-β is one of the most important stimulators of extracellular matrix production [28]. As such, we evaluated gene expression of ECM coding genes, including fibronectin (FN), collagen1A (COL1A), and collagen 3A (COL3A) in HAO at 10-DPI with rHPh-TX H5N1-Venus or pH1N1-mCherry. We observed a ∼15-fold increase in FN expression caused by HPh-TX H5N1-Venus infection and a ∼2.5-fold increase in pH1N1-mCherry-infected HAO (**Fig. 5D**). Similarly, COL1A and COL3A increased ∼4-fold by HPh-TX H5N1-Venus infection and ∼2-fold during pH1N1-mCherry infection (**Figs. 5E and 5F**). These increase in the ECM gene expression was combined with upregulation of tissue remodeling matrix metalloproteinase MMP2 (**Fig. 5G**) and MMP9 (**Fig. 5H**) enzymes that were upregulated ∼15-fold after rHPh-TX H5N1-Venus infection *vs.* ∼2-fold upregulation in pH1N1-mCherry-infected HAO. These results suggest that rHPh-TX H5N1 infection induces a more pronounced EMT and ECM remodeling than pH1N1, as evidenced by the significantly higher upregulation of TGF-β and ECM-associated genes. The robust increase in FN, COL1A, and COL3A expression, along with the marked upregulation ofMMP2 and MMP9, highlights the potential for H5N1 to drive a profibrotic response in infected epithelial tissues. Given the well-established role of TGF-β in promoting fibrosis [19], these findings suggest that H5N1 infection may contribute to long-term lung remodeling and fibrotic pathology through sustained EMT and ECM deposition. Understanding these mechanisms could provide insight into the differential pathogenicity of IAV strains and inform targeted therapeutic strategies to mitigate fibrosis-associated complications.

### ROCK1 activity is required for fibroblast-like cell foci formation caused by HPh-TX H5N1 infection in HAO

Our data suggest that H5N1 infection caused higher HAO remodeling and upregulation of the fibrotic cytokines and ECM deposition compared to pH1N1. These findings align with the distinct pathogenicity of H5N1 and long-term consequences such as pneumonia, lung fibrosis, and respiratory failure [29, 30]. Unlike pH1N1, which typically causes self-limiting respiratory illness in immunocompetent individuals, H5N1 infection is associated with severe lung injury, prolonged inflammation, and a higher likelihood of developing post-viral fibrosis [15, 24]. As such, we sought a therapeutic approach to tackle the H5N1 pathogenesis. The Rho-associated coiled-coil containing protein kinases (ROCK1 and ROCK2) signaling pathway plays a central role in regulating cytoskeletal dynamics, EMT, and ECM remodeling that are critically involved in fibrogenesis [19]. We investigated the role of ROCK isoforms in HAO fibrosis after infection with a HPh-TX H5N1 natural isolate. Using virus natural isolate better reflects the genetic makeup of the circulating strain, making findings more translatable to clinical settings. To this end, we evaluated the effects of selective ROCK1 (GSK269962A) and ROCK2 (Belumosudil/KD025) inhibitors on the fibroblast-like cell foci formation after infecting HAO (MOI 0.001) with HPh-TX H5N1 for 10 days. Infected HAO were treated with a non-toxic dose of ROCK1 and ROCK2 inhibitors (5 µM) for 5 days, starting from 5-DPI to 10-DPI, the period of fibroblast-like cell foci formation (**Fig. 4A**). On 10-DPI, the tissue was collected, fixed, and immunostained with a FITC-conjugated antibody against α-SMA (**Fig. 6A**). ROCK1 inhibitor-treated HAO showed reduced α-SMA expression, with no detectable fibroblast-like foci formation. In contrast, ROCK2 inhibitor-treated HAO exhibited higher α-SMA expression and retained fibroblast-like foci (**Fig. 6A**). We also measured the gene expression of the extracellular matrix protein, which showed a significant reduction of FN, COL1A, and COL3A in ROCK1-treated HAO compared to mock-treated or ROCK2-treated tissues (**Figs. 6B-D**). We also observed a significant decrease in extracellular matrix-modulating enzyme MMP2 (**Fig. 6E**) and MMP9 (**Fig. 6F**) in ROCK1-treated HAO *vs.* mock-treated or ROCK2-treated tissues. These results suggest that inhibition of ROCK1, but not ROCK2, effectively suppresses fibroblast-like foci formation and ECM remodeling in HPh-TX H5N1-infected HAO. The reduction in α-SMA expression and the downregulation of FN, COL1A, and COL3A in ROCK1 inhibitor-treated HAO indicate that ROCK1 plays a pivotal role in driving EMT and fibrosis. Furthermore, ROCK1 inhibition significantly reduced MMP2 and MMP9 expression, suggesting reduced ECM degradation and remodeling. In contrast, ROCK2 inhibition failed to prevent fibroblast-like foci formation and was associated with sustained ECM protein expression, highlighting the distinct roles of ROCK isoforms in fibrosis.

**Figure 6:**
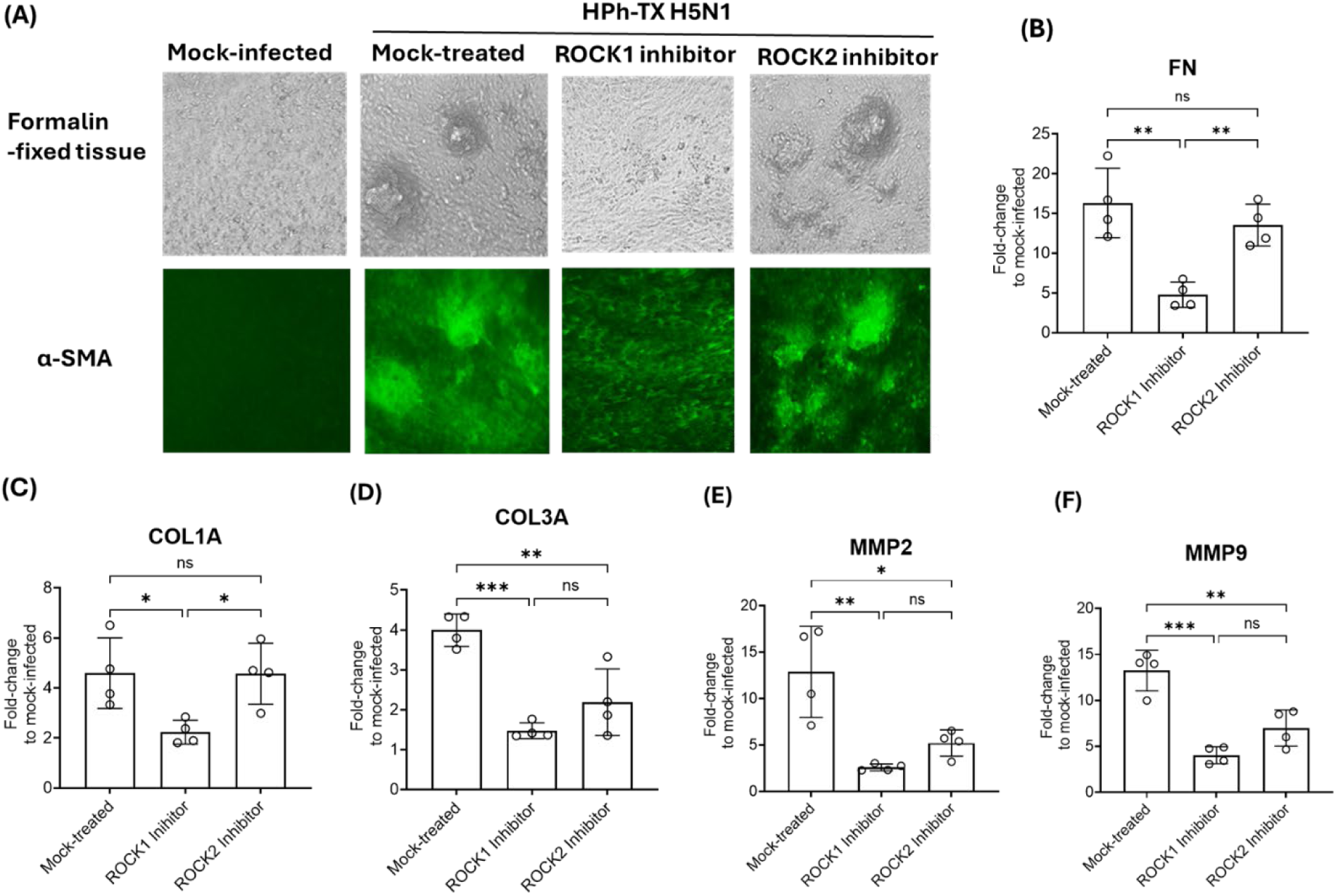
Inhibition of ROCK1 impaired fibroblast-like cell foci formation after infection of HAO with HPh-TX H5N1. **(A)** HAO tissue was infected (MOI 0.001) with HPh-TX H5N1 and treated with a non-toxic dose (5 μM) of ROCK1 (GSK269962) or ROCK2 (KD025, Belumosudil) inhibitors from 5-DPI to 10-DPI. HAO tissue was fixed and stained with α-SMA (green) at 10-DPI. **(B-D)** Gene expression of extracellular matrix proteins fibronectin (FN), collagen 1A (COL1A), and collagen 3A (COL3A). (**E**, **F**) Gene expression of tissue remodeling MMP2 and MMP9. Data were analyzed using One-way ANOVA followed by a post-test comparison. The significant differences are indicated (* = *p* < 0.05, ** = *p* < 0.01, *** = *p* < 0.001; non-significant = ns).

## DISCUSSION

The unpredictable evolution of IAVs makes future pandemics inevitable, highlighting the urgent need for preparedness to mitigate health, social, and economic impacts. H5N1 cases were first described in Hong Kong in 1997 when birds died from an outbreak of the disease, propagating to humans later in the same year [31, 32]. Since the resurgence of H5N1 in 2002–2003, global efforts have been intensified to prevent its spread. Recently, a multistate outbreak of H5N1 in dairy cattle was reported in the Spring of 2024 in the United States, resulting in several human infections and 1 death[33]. This outbreak underscores the risk of cross-species transmission as the virus has spread within poultry, cattle, and humans, posing a significant public health threat [34].

Infections with H5N1 cause severe pneumonia that can progress to lung fibrosis and respiratory failure [29, 30]. While current FDA-approved antivirals reduce viral loads [35], these antivirals are often ineffective in preventing lung injury and fibrosis, underscoring the need for novel therapeutic strategies targeting host responses. To better understand viral replication and pathogenesis, we employed HAO, a physiologically relevant model that recapitulates key aspects of the human lung environment.

The data of this study suggest that rHPh-TX H5N1 replicates efficiently in HAO and can cause severe infection in human airway epithelial tissues. The high levels of infectious virus particles in secreted mucus could be a source of the virus in aerosols from infected subjects, increasing the likelihood of person-to-person virus transmission. Our findings demonstrate that rHPh-TX H5N1 efficiently replicates in HAO, eliciting a strong IFN response. Typically, H5N1 infection activates the RIG-I/MAVS signaling pathway, leading to IRF3 and NF-κB activation, which drives IFN-β production and ISG15 expression [36–38]. Our data show that rHPh-TX H5N1 induces IRF3 and NF-κB expression, which increase IFN-β and ISG15 levels. While ISG15 contributes to antiviral defense, its dual role in immune regulation can either enhance or suppress IRF3/NF-κB signaling, potentially influencing disease severity. In severe cases, H5N1 evades IFN pathways, leading to excessive NF-κB-driven inflammation (cytokine storm) and impaired IRF3-mediated responses, exacerbating lung damage [39–41]. Cytokine profiling of infected HAO revealed that rHPh-TX H5N1 induced elevated IL-6, IL-1β, TNF, CCL5, and IP-10 (CXCL10) levels. While these cytokines aid in viral clearance, prolonged inflammation may contribute to tissue injury and fibrosis (e.g. cytokine storm-induced lung injury).

Given the robust inflammatory response induced by rHPh-TX H5N1, we investigated its role in airway fibrogenesis during prolonged infection with rHPh-TX H5N1 compared to seasonal pH1N1. Our results indicate that prolonged infection of HAO with rHPh-TX H5N1 and pH1N1 induced fibroblast-like cells surrounding the infected area, associated with high cytokine response and α-SMA expression, indicating fibroblast-to-myofibroblast differentiation. It has been reported that H1N1 infections can rapidly progress to ARDS and contribute to pulmonary fibrosis [42]. Furthermore, severe IAV infections are associated with higher levels of TGF-β, indicating a relationship between disease severity and IAV-induced pulmonary fibrosis [43]. This agrees with our findings showing that the rHPh-TX H5N1 infection induces higher TGF-β and ECM-associated gene expression than pH1N1. As such, the increased fibroblast activity in post-inflammatory repair pathways, with a pivotal role played by TGF-β and ECM, seems to be linked to IAV-induced pulmonary fibrosis.

We hypothesize that epithelial cells underwent the EMT process that plays a prominent role in fibrogenesis, where epithelial cells lose cell-cell attachment, polarity, and epithelial-specific markers, undergo cytoskeletal remodeling, and gain a mesenchymal phenotype [44]. Our results indicate that rHPh-TX H5N1 infection induces more pronounced EMT and ECM remodeling than pH1N1, as evidenced by the significantly higher upregulation of TGF-β and ECM-associated genes. We also observed the spindle shape of fibroblasts with intensive expression of α-SMA. In this regard, activated fibroblasts are described as a spindle or stellate morphology with intracytoplasmic stress fibers, a contractile phenotype, expression of various mesenchymal immunocytochemical markers (including, most reliably, α-SMA, and collagen production [25].

We also observed that at 10-DPI, pro-inflammatory (TNF, IL-6, IL-8, IL-1β) and pro-fibrotic (TGF-β) mediators were upregulated, along with ECM components (fibronectin, collagen 1A, and collagen 3A) and matrix metalloproteinases MMP2 and MMP9. Importantly, the upregulation of fibronectin expression allows microbes to adhere to epithelial surfaces and contributes to the virulence of secondary bacterial infections [45, 46]. Collectively, these factors contribute to a feedback loop of persistent lung damage and fibrosis [24], emphasizing the need for targeted interventions to disrupt these pathways. Our data showed that the ROCK pathway affects fibroblast foci formation and ECM deposition. This pathway is a key regulator of fibrosis and exists in two isoforms (ROCK1/ROCK2) with distinct roles in tissue remodeling [19]. Our results indicate that the inhibition of the ROCK1 isoform significantly reduced fibroblast foci formation and ECM deposition, whereas ROCK2 inhibition had a lesser effect. This suggests a dominant role for ROCK1 in airway fibrosis during H5N1 infection. Previous studies described the capacity of antifibrotic agents such as pirfenidone and nintedanib, FDA-approved to treat idiopathic pulmonary fibrosis (IPF), to reduce fibrosis through inhibiting key cytokines, including TGF-β and vascular endothelial growth factor (VEGF) [47, 48]. Thus, unraveling the mechanisms of the ROCK signaling pathway in IAV-induced fibrogenesis may be essential for developing effective strategies to prevent and treat viral-induced pulmonary fibrosis, ultimately mitigating its long-term impact on respiratory health.

While our study provides insights into H5N1 pathogenesis and triggering fibrogenesis, it also has limitations. Though physiologically relevant, the HAO model does not fully recapitulate *in vivo* immune responses or systemic factors influencing disease progression. Thus, future studies using *in vivo* models are needed to validate our findings. Additionally, the potential therapeutic efficacy of ROCK1 inhibitors should be tested in preclinical models to assess their impact on IAV-induced lung injury and fibrosis. Future studies should also explore the molecular mechanisms linking IFN signaling, inflammation, and fibrosis to identify additional therapeutic targets.

In conclusion, our findings highlight the efficient replication and immune response activation of H5N1 in HAO. H5N1 induces a robust inflammatory and fibrotic response, driven by NF-κB and TGF-β signaling, contributing to airway remodeling and fibrosis. Targeting ROCK1 represents a potential strategy to mitigate IAV-induced lung fibrosis. These insights emphasize the importance of developing host-targeted therapies to prevent severe lung complications associated with influenza infections.

## AUTHORS CONTRIBUTIONS

Conceptualization: H.R. and L.M-S.; Methodology: H.R., A.M., M.B., C.Y., R.S.B., and A.A-G.; Data collection and interpretation: H.R. and L.M-S.; Funding acquisition and resources: H.R. and L.M-S.; Writing, review, and editing: H.R., A.N.J., J.B.T., and L.M-S.; all authors have read and agreed to the published version of the manuscript.

## ACKNOWLEDGMENTS

This work was supported by a grant from the American Lung Association (ALA) to L.M-S. and the Texas Biomed Forum Award to H.R.

## Notes

### Competing Interest Statement

The authors have declared no competing interest.

